# Microsatellite primer development for invasive perennial herb, *Gypsophila paniculata* (Caryophyllaceae)

**DOI:** 10.1101/401398

**Authors:** Hailee B. Leimbach-Maus, Syndell R. Parks, Charlyn G. Partridge

**Author notes:** Email addresses: HBLM, SRP, CGP. Manuscript received _____; revision accepted_____. The authors thank the Grand Valley State University Center for Scholarly and Creative Excellence Catalyst Grant (C.G.P) and the Environmental Protection Agency – Great Lakes Restoration Initiative Grant (C.G.P., Grant #00E01934) for financial support. The authors also thank E.K. Rice for help in field sampling, and M. Kienitz for help in the laboratory.

## Abstract

- *Premise of the study: Gypsophila paniculata* L. (baby’s breath) is an herbaceous perennial that has invaded much of northern and western United States and Canada, outcompeting and crowding out native and endemic species. Microsatellite primers were developed to analyze the genetic structure of invasive populations.
- *Methods and Results:* We have identified 16 polymorphic nuclear microsatellite loci for *G. paniculata* out of 73 loci that successfully amplified from a primer library created using Illumina sequencing technology. The developed primers amplified microsatellite loci in 3 invasive populations in Michigan. Primers amplified di-, tri-, and tetra-nucleotide repeats.
- *Conclusions:* These markers will be useful in characterizing the genetic structure of invasive populations throughout North America to aid targeted management efforts, and in native Eurasian populations to better understand invasion history. Five of these developed primers also amplified in *G. elegans*.

## INTRODUCTION

The herbaceous perennial forb, *Gypsophila paniculata* L., was introduced to North America in the late 1800’s (Darwent and Coupland, 1966). Invasive populations have since been documented throughout the northern and western United States and Canada, specifically in agriculture fields, rangeland, roadsides, and sandy coastlines along the Great Lakes (Darwent, 1975; Emery and Doran, 2013). Despite its wide invasive range, little information exists on how populations throughout North America are related or spreading. Due to its aggressive invasion, negative impacts on native biota (Emery and Doran, 2013), and a lack of data regarding its spread, it is important to develop molecular markers that can characterize the genetic structure of invasive populations of *G. paniculata*. These markers will be directly used to investigate invasions within the Lake Michigan coastal dune system where an 1,800-acre infestation occurs (TNC, 2013). These markers and optimized protocols can be used to characterize populations of *G. paniculata* throughout its invasive and native ranges to further assess its invasion history and spread.

Calistri et al. (2014) examined the genetic relationship of five *Gypsophila spp*. (including *G. paniculata*) within their native range and 13 commercial hybrid strains using a combination of amplified fragment length polymorphisms (AFLPs), inter simple sequence repeats (ISSRs), target region amplification polymorphism (TRAP), and universal chloroplast simple sequence repeats (cpSSRs). However, the majority of these markers are dominant and thus do not fully distinguish between homozygotes and heterozygotes, a characteristic that would allow for fine-scale population genetic analyses (Freeland et al., 2011). Thus, the development of microsatellite markers for *G. paniculata* is necessary to adequately characterize invasive populations throughout North America.

## METHODS AND RESULTS

### Microsatellite Library Development, Assembly and Identification

Adventitious buds growing from the caudex of five *G. paniculata* plants were collected from Sleeping Bear Dunes National Lakeshore (hereafter Sleeping Bear Dunes or SBDNL) in 2015 to develop the microsatellite library. Tissue was stored in indicator silica until DNA extraction. Genomic DNA was extracted using DNeasy plant mini kits (QIAGEN, Hilden, Germany), with modifications including extra wash steps with AW2 buffer. Extracted DNA was run through Zymo OneStep PCR Inhibitor Removal Columns twice (Irvine, California, USA), and checked using a Thermo Fisher Scientific Nanodrop 2000 (Waltham, Massachusetts, USA). For microsatellite library development, each sample was diluted to 50 ng/µL and submitted to Cornell University, Department of Ecology and Evolutionary Biology. Libraries were then submitted to the Sequencing and Genotyping Facility at the Cornell Life Science Core Laboratory Center for sequencing using a 2×250 paired-end format on an Illumina MiSeq (Appendix S1). Raw sequence files for the microsatellite library have been deposited to NCBI’s Short Read Archive (Bioproject No: PRJNA431197). A total of 58,907 contigs containing microsatellite loci were obtained. Msatcommander (v 1.0.3) (Faircloth, 2008) identified 3,892 potentially unique primers that yielded products of 150-450 bp, had a GC content between 30-70%, and that had a T_m_ between 58-62°C, with an optimum of 60°C (Appendix S2).

### Primer Optimization

Prior to PCR optimization, contigs containing potential primers were aligned using ClustalOmega (Sievers et al., 2011) to ensure they were targeting unique microsatellite regions. We tested 107 primer pairs that consisted of either tetrameric, trimeric, or dimeric motifs, and yielded products between 150-300 bp. Of these, 73 successfully amplified, and 16 were determined to be polymorphic and easily scorable (Appendix S3). DNA from leaf tissue collected in 2016 from three populations (Zetterberg Preserve, SBDNL, Petoskey State Park) along eastern Lake Michigan was used for primer optimization (see Table 2 for population geographic coordinates). A minimum of 30 tissue samples were collected from each population. Tissue storage and DNA extraction methods are the same as previously stated.

PCR reactions consisted of 1x KCl buffer (Thermo Fisher, Waltham, Massachusetts, USA), 2.0-2.5 mM MgCl_2_ (Table 1^b^) (Thermo Fisher, Waltham, Massachusetts, USA), 300 µM dNTP (New England BioLabs, Ipswich, Massachusetts, USA), 0.08 mg/mL BSA (Thermo Fisher, Waltham, Massachusetts, USA), 0.4 µM forward primer fluorescently labeled with either FAM, VIC, NED, or PET (Applied Biosystems, Foster City, California, USA), 0.4 µM reverse primer (Integrated DNA Technologies, Coralville, Iowa, USA), 0.25 units of *Taq* polymerase (Thermo Fisher, Waltham, Massachusetts, USA), and a minimum of 50 ng DNA template, all in a 10.0 µL reaction volume. The thermal cycle profile consisted of 94°C for 5 minutes, 35 cycles of 94°C for 1 minute, primer-specific annealing temperature (Table 1) for 1 min, 72°C for 1 min, and a final elongation step of 72°C for 10 minutes. Successful amplification was determined by visualizing the amplicons on a 2% agarose gel stained with ethidium bromide. Fragment analysis of the amplicons was performed on an ABI3130xl Genetic Analyzer (Applied Biosystems, Foster City, California, USA).

**Table 1.**
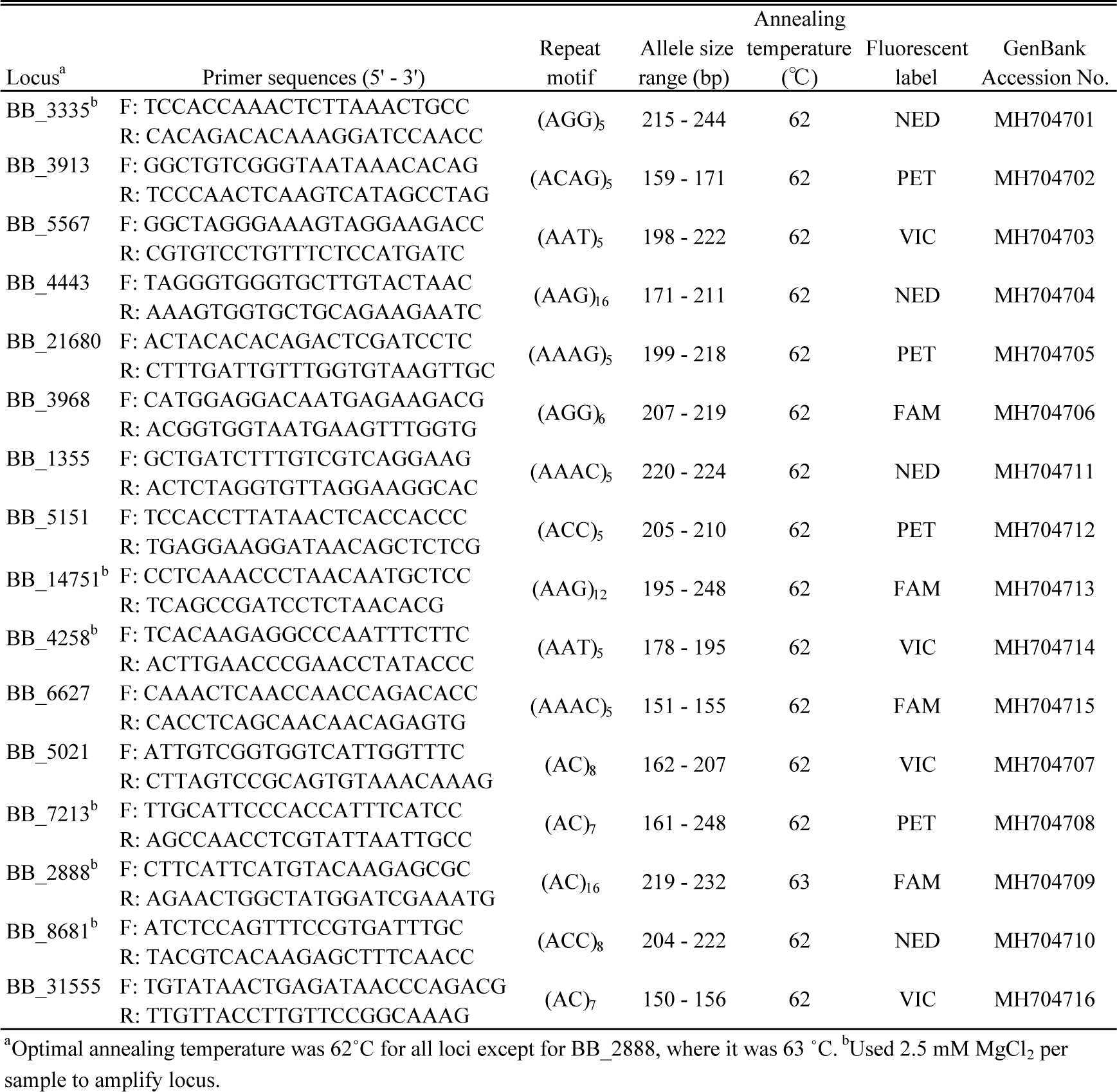
Characteristics of 16 nuclear microsatellite loci developed for *Gypsophila paniculata.*

**Table 2.** Genetic properties of the 16 developed nuclear microsatellite loci for 3 populations of *Gypsophila paniculata.*

| Locus | Zetterberg Preserve ($n = 30$) | | | | | Sleeping Bear Dunes ($n = 33$) | | | | | Petoskey State Park ($n = 30$) | | | | |
| --- | --- | --- | --- | --- | --- | --- | --- | --- | --- | --- | --- | --- | --- | --- | --- |
| | $A$ | $H_0^a$ | $H_E$ | $F_{IS}$ | $r$ | $A$ | $H_0^a$ | $H_E$ | $F_{IS}$ | $r$ | $A$ | $H_0^a$ | $H_E$ | $F_{IS}$ | $r$ |
| BB_3335 | 8 | 0.70 | 0.82 | 0.09 | 0.07 | 7 | 0.80 | 0.80 | -0.01 | 0.00 | 3 | 0.50 | 0.56 | 0.02 | 0.05 |
| BB_3913 | 4 | 0.74 | 0.68 | -0.04 | -0.06 | 4 | 0.59 | 0.61 | 0.05 | 0.00 | 2 | 0.13 | 0.13 | -0.06 | -0.06 |
| BB_5567 | 4 | 0.63 | 0.60 | -0.04 | -0.04 | 4 | 0.63 | 0.63 | 0.01 | -0.01 | 3 | 0.53 | 0.62 | 0.14 | 0.05 |
| BB_4443 | 10 | 0.60 | 0.64 | 0.03 | 0.03 | 8 | 0.76 | 0.77 | -0.02 | 0.00 | 5 | 0.80 | 0.71 | -0.06 | -0.09 |
| BB_21680 | 4 | 0.53 | 0.56 | 0.01 | 0.02 | 3 | 0.52 | 0.56 | 0.02 | 0.05 | 4 | 0.67* | 0.58 | 0.23 | -0.10 |
| BB_3968 | 3 | 0.50 | 0.42 | -0.14 | -0.11 | 4 | 0.35* | 0.49 | 0.15 | 0.13 | 2 | 0.04 | 0.04 | — | 0.28 |
| BB_5021 | 5 | 0.77 | 0.73 | 0.00 | -0.05 | 5 | 0.54 | 0.60 | 0.15 | 0.02 | 1 | — | — | — | 0.30 |
| BB_7213 | 5 | 0.63 | 0.61 | -0.05 | -0.02 | 3 | 0.59 | 0.58 | -0.13 | 0.04 | 2 | 0.37 | 0.38 | 0.04 | 0.01 |
| BB_2888 | 6 | 0.90 | 0.80 | -0.11 | -0.07 | 6 | 0.71 | 0.82 | 0.20 | 0.06 | 2 | 0.57 | 0.50 | -0.13 | -0.08 |
| BB_8681 | 4 | 0.47 | 0.51 | 0.23 | 0.01 | 4 | 0.38 | 0.36 | -0.07 | -0.05 | 3 | 0.41 | 0.53 | 0.28 | 0.17 |
| BB_1355 | 2 | 0.17 | 0.26 | 0.37 | 0.13 | 2 | 0.14 | 0.13 | -0.07 | -0.08 | 2 | 0.37 | 0.51 | 0.29 | 0.13 |
| BB_5151 | 2 | 0.20 | 0.18 | -0.10 | -0.11 | 2 | 0.49 | 0.51 | 0.11 | 0.00 | 2 | 0.03 | 0.03 | — | -0.02 |
| BB_14751 | 6 | 0.79 | 0.71 | -0.06 | -0.07 | 6 | 0.76 | 0.76 | -0.02 | -0.01 | 3 | 0.20 | 0.29 | 0.21 | 0.12 |
| BB_4258 | 3 | 0.40 | 0.33 | -0.11 | -0.22 | 1 | 0.00 | 0.00 | — | 0.00 | 1 | — | — | — | 0.00 |
| BB_6627 | 2 | 0.30 | 0.26 | -0.16 | -0.16 | 2 | 0.46 | 0.49 | -0.03 | 0.04 | 1 | — | — | — | 0.00 |
| BB_31555 | 4 | 0.60 | 0.67 | 0.07 | 0.05 | 4 | 0.46 | 0.59 | 0.11 | 0.12 | 2 | 0.23 | 0.26 | 0.10 | 0.04 |

Note: $n$ = number of individuals sampled; $A$ = number of alleles; $H_0$ = observed heterozygosity; $H_E$ = expected heterozygosity; $F_{IS}$ = fixation index; $r$ = null allele frequency; — = data not available because only 1 allele present at locus or because $A > 1$ is due to rare allele.
Data based on 3 populations with the following geographic coordinates: Zetterberg Preserve = 44.68231, -86.25322; Sleeping Bear Dunes = 44.87372, -86.06170; Petoskey State Park = 44.40859, -84.91238. All 3 populations are located in the northwest region of Michigan's lower peninsula.
\*Significant deviation from Hardy-Weinberg equilibrium: \* $P < 0.05$ .

### Microsatellite marker data analysis

Alleles were scored using Genemapper v5 (Applied Biosystems, Foster City, California, USA), and Micro-Checker v2.2.3 (Van Oosterhout et al., 2004; Van Oosterhout et al., 2006) was used to identify null alleles and potential scoring errors from stuttering or large allele dropout. There was no significant evidence of null alleles (p > 0.05) in the Zetterberg Preserve and SBDNL populations. However, null alleles were suggested for loci BB_3968, BB_5021, and BB_8681 in the Petoskey State Park population (Table 2). Homozygote excess for Petoskey State Park is not surprising, given this population’s reduced number of alleles at each locus and small comparative population size. We characterized genetic diversity by examining the number of alleles, and expected and observed heterozygosity for each locus averaged over each population (Table 2) using the package STRATAG in the R statistical program (Archer et al., 2016). The number of observed alleles ranged from 1 – 10. Some loci were monomorphic for one population, but polymorphic when analysis included all populations (e.g., BB_4258).

The Zetterberg Preserve population displayed slightly higher heterozygosity values than Sleeping Bear Dunes, but the Petoskey State Park population had much lower heterozygosity values in comparison. A probability test for Hardy-Weinberg Equilibrium (HWE), calculation of the fixation index (F_IS_), and linkage disequilibrium were performed in GENEPOP 4.2 (Raymond and Rousset, 1995; Rousset, 2008). The default parameters for Markov Chain Monte Carlo (MCMC) iterations were used to calculate HWE. All loci were in HWE except locus BB_3968 for SBDNL, and locus BB_21680 for Petoskey State Park (Table 2). The F_IS_ estimates were calculated using the probability model following Robertson and Hill (1984). Statistical tests for genetic linkage disequilibrium were performed using the log likelihood ratio statistic (G-test) and MCMC algorithm by Raymond and Rousset (1995). Two pairs of loci were significantly out of linkage disequilibrium (p < 0.05) for both Zetterberg Preserve and Sleeping Bear Dunes: BB_5021 and BB_2888, and BB_3913 and BB_1355. Out of 16 loci, five successfully amplified in a related species *G. elegans* (BB_4443, BB_4258, BB_7213, BB_5151, BB_1355) (Table 3).

**Table 3.** Results of cross - amplification of microsatellite loci isolated from *Gypsophila paniculata* and tested in 12 *G. elegans* individuals. ^a^

|  |  |  |  |  |  |  |  |  |  |  |  |  |  |
| --- | --- | --- | --- | --- | --- | --- | --- | --- | --- | --- | --- | --- | --- |
| Successful amplification (X) in each *G. elegans* individuals (*n* = 12). |  |  |  |  |  |  |  |  |  |  |  |  |  |
| Locus | Allele size<br>range (bp) | GYEL01 | GYEL02 | GYEL03 | GYEL04 | GYEL07 | GYEL08 | GYEL09 | GYEL12 | GYEL13 | GYEL15 | GYEL18 | GYEL19 |
| BB_3335 |  |  |  |  |  |  |  |  |  |  |  |  |  |
| BB_3913 |  |  |  |  |  |  |  |  |  |  |  |  |  |
| BB_5567 |  |  |  |  |  |  |  |  |  |  |  |  |  |
| BB_4443 | 152 | X | X | X | X | X | X | X | X | X | X | X | X |
| BB_21680 |  |  |  |  |  |  |  |  |  |  |  |  |  |
| BB_3968 |  |  |  |  |  |  |  |  |  |  |  |  |  |
| BB_1355 | 209 - 222 | X | X | X | X | X | X | X | X | X | X | X | X |
| BB_5151 | 202 - 205 | X | X | X | X | X | X | X | X | X | X | X | X |
| BB_14751 |  |  |  |  |  |  |  |  |  |  |  |  |  |
| BB_4258 | 160 - 169 | X | X | X | X | X | X | X | X | X | X | X | X |
| BB_6627 |  |  |  |  |  |  |  |  |  |  |  |  |  |
| BB_5021 |  |  |  |  |  |  |  |  |  |  |  |  |  |
| BB_7213 | 161 - 171 | X | X | X | X | X | X | X | X | X | X | X | X |
| BB_2888 |  |  |  |  |  |  |  |  |  |  |  |  |  |
| BB_8681 |  |  |  |  |  |  |  |  |  |  |  |  |  |
| BB_31555 |  |  |  |  |  |  |  |  |  |  |  |  |  |

<sup>a</sup>*G. elegans* tissue sourced from individuals grown in greenhouse at AWRI-GVSU facilities.

## CONCLUSIONS

The 16 microsatellite primers developed for *G. paniculata* provide a tool for estimating genetic diversity and structure of invasive populations, which will aid in understanding its invasion history, identifying source populations, and examining dispersal patterns. Though we developed these markers to study the Lake Michigan dune system invasion, it is invasive throughout North America. With these markers, we can begin to understand the invasion of *G. paniculata* in North America in order to improve management efforts and prevent the further spread of this species.

## DATA ACCESSIBILITY

A summary of the microsatellite library development and sequence analysis protocols (unpublished data) provided to us by Cornell University, Department of Ecology and Evolutionary Biology are in Appendix S1. Fasta sequences for the 16 microsatellite primers developed here are in Appendix S5. The fasta file listing all identified contigs containing microsatellite regions are in Appendix S4. Potential primer pairs for the identified microsatellite-containing contigs are in Appendix S2. The 107 *G. paniculata* – specific primer pairs tested during primer optimization are in Appendix S3. Raw sequence files for the microsatellite library have been deposited to NCBI’s Short Read Archive (Bioproject No: PRJNA431197) and microsatellite sequences have been deposited to GenBank (Table 1). Voucher specimen for each population have been deposited into the Grand Valley State University Herbarium (GVSC), Grand Valley State University Department of Biology, Allendale, MI, USA. (Appendix S6).

